# Cultural norms of exogamy and mobility shape hunter-gatherer genetic evolution

**DOI:** 10.64898/2026.03.10.710787

**Authors:** Cecilia Padilla-Iglesias, Dieudrice Nganga, Eustache Amboulou, Juliette Ruf, Pascale Gerbault, Mylène Docquier, Suspense Ifo Averti, Lucio Vinicius, Andrea Manica, Andrea Migliano

**Author notes:** These authors contributed equally as senior authors (Andrea Manica, Andrea Migliano).

## Abstract

Through a hunting and gathering lifestyle, humans have managed to thrive across all terrestrial ecosystems. A key adaptive feature enabling this ecological success is the ability of hunter-gatherer societies to maintain high levels of genetic diversity despite ecological and demographic shocks, but the mechanisms underlying this resilience are poorly understood. Here we integrate genomic, demographic, mobility and ethnographic data from two Central African hunter-gatherer populations to show that genetic diversity emerges from interacting effects of population size, mobility and cultural norms governing marriage. We first demonstrate direct selection against background homozygosity: even modest increases in runs of homozygosity, in the near absence of close-kin marriage, are associated with reduced reproductive success. Despite regional differences in effective population size, clustering of relatives, sedentism and exogamy rules, overall levels of homozygosity are similarly low in both populations. These shared genetic outcomes are achieved through distinct strategies: in one region, strict exogamy combined with high lifetime mobility limits local relatedness, whereas in the other, more relaxed exogamy norms are offset by increased male mate-search distances that reduce offspring homozygosity. Together, our results show that human populations flexibly adjust mobility and social norms to demographic constraints to preserve genetic diversity and avoid fitness costs, revealing culture as a central component of human adaptation.

## Introduction

For most of our evolutionary history, humans lived as hunter-gatherers^1^. This lifestyle has proven remarkably resilient, thriving across every terrestrial ecosystem on the planet, and enduring major climatic shifts throughout the Pleistocene and Holocene. Research in both contemporary and ancient DNA has revealed that part of what has made this lifestyle resilient is the ability of hunter-gatherer populations to maintain high levels of genetic diversity (low inbreeding and high levels of heterozygosity) despite low population densities and census population sizes^2–7^.

At the same time, ethnographic and anthropological research has pointed to hunter-gatherer social structures - characterised by fluid residential groups, extensive inter-camp ties, and networks of bilateral kinship - as key for ensuring such high levels of genetic diversity^2,4,8–12^. The two main interconnected processes linking hunter-gatherer social structures and genetic diversity are mobility and marriage practices, which in humans are the main institution regulating reproduction. In most animal societies, individuals of one or both sexes disperse to find mates, with important consequences for the maintenance of genetic diversity. In humans, mobility additionally structures cultural exchange and social alliances, including marital partner choice^13–16^. At the same time, cultural norms governing marriage practices regulate which partners are suitable, from where they may be drawn, and where couples will reside after marriage - thereby also shaping mobility^17–20^.

The marital exchange of mates among kin groups (reciprocal exogamy) and accompanying networks of economic exchange have long been considered central to the organisation of hunter-gatherer societies^19–21^. Clan-based exogamy systems, present in many hunter-gatherer societies, often require individuals to travel to distant camps to find suitable marriage partners, thereby creating bilateral kinship ties that extend social networks across space^22–25^. Ancient DNA evidence suggests that such exogamous marriage practices operated in the Late Pleistocene in Europe and Africa, where patterns of genetic relatedness indicate systematic avoidance of close kin mating and extensive gene flow between sites separated by over 100km^4,26^.

However, hunter-gatherer social structures and demographic contexts are highly variable, both across space and through time^1,27–29^. Population densities, group sizes, and the availability of potential mates differ substantially among populations and fluctuate with environmental conditions. Whether and how hunter-gatherer societies adapt mobility, marriage and cultural practices to ensure high levels of genetic diversity and to mitigate negative fitness effects of inbreeding remains poorly understood.

Inbreeding, the mating between genetically closely related individuals, arises either through consanguineous marriages or the fixation of genetic variants due to genetic drift. It therefore leads to an increase in homozygosity in the offspring, and is often quantified by measuring the fraction of an individual’s genome in runs of homozygosity (F_ROH_)^2,30,31^. Elevated F_ROH_ has been associated with decreased height, adverse cardiometabolic phenotypes, and elevated incidence of recessive disorders^32–35^, and in several animal species, increased homozygosity is linked to higher mortality^36–38^. This is because long runs of homozygosity tend to expose deleterious variants in homozygote states that would otherwise not be expressed in heterozygotes^39^.

In our own species, studies using pedigree data have shown that increased parental relatedness elevates childhood mortality and can decrease the total number of surviving offspring^40–42^, yet studies linking genomic estimates of inbreeding with fitness consequences are lacking. One exception was the study by Swinford et al.^43^ that provided the first direct evidence linking homozygosity to negative fitness consequences among the Himba of Namibia. However, this is an endogamous population with exceptionally high background levels of consanguinity, which may not be representative of the hunter-gatherer populations that characterised our species for most of its history, where consanguineous matings are extremely rare and exogamy norms commonplace^44–48^.

At the same time, empirical data quantifying the distances people travel to find marriage partners in hunter-gatherer societies are also limited^1,25,49^. Early theoretical models proposed that hunter-gatherer range sizes should be jointly determined by subsistence and reproductive concerns: minimum ranges set by resource requirements, and maximum ranges by the need to access an effective mating pool^50–52^. MacDonald and Hewlett^25^ provided some empirical support for these ideas, showing across 5 forager societies that lower population density was associated with greater mating distances—the average distance between birthplaces of couples. Among the Aka of the Central African Republic, they found that as sedentarisation increased, population density decreased logarithmically, mating distances increased linearly, and that this relationship held exclusively for men, suggesting that male range expansion is partly driven by mate-seeking. Cashdan^53^ observed how when the G//ana of the Kalahari transitioned from mobile to more sedentary, their mating distance decreased. These findings imply that sedentarisation should result in higher levels of inbreeding, which in turn could have negative fitness consequences. However, no study has yet traced the full process from local demographic conditions through spatial behaviours and cultural norms to social structures, genetic outcomes and, ultimately, fitness consequences that could be evolutionarily relevant (Figure 1; see^54^).

**Figure 1.**
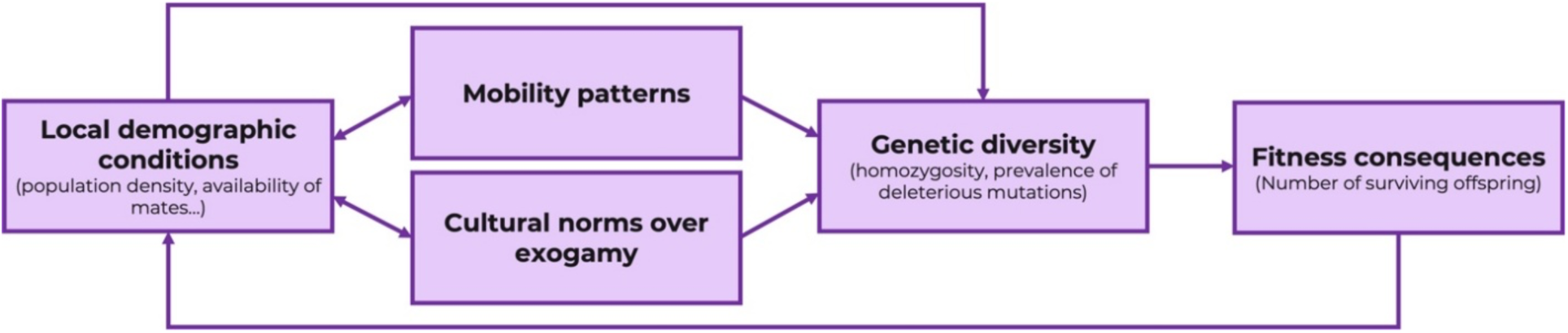
Hypothesized relationships between local demographic conditions, mobility, cultural norms, social structures, genetic diversity patterns and fitness consequences.

Central African hunter-gatherers offer an exceptional opportunity to investigate these relationships. These populations represent one of the most divergent human genetic lineages, having separated from other African populations as far back as 120,000–250,000 years ago^26,55,56^, and they are thought to descend from hunter-gatherers who inhabited the Congo Basin throughout the late Pleistocene. The BaYaka of the Republic of Congo are organized into named clans that extend beyond single territories and provide access to multiple stretches of forest. Previous ethnographic work has reported these clans to be patrilineally inherited and exogamous across all Central African hunter-gatherer populations^45,57,58^. However, our fieldwork revealed striking variation in marriage rules between two regions: in Macao, all respondents to our demographic questionnaire (N=185) stated that individuals are prohibited from marrying members of both their paternal and maternal clans, whereas in Minganga, all respondents (N=58) reported that only paternal clan exogamy is enforced.

Previous research (see also Figure S1) documented differences in mobility patterns between these regions, with individuals in Macao having showing larger lifetime spatial ranges^29^. It also showed differences in reasons for travel, and the magnitude of sex differences in movement, alongside potential fitness consequences of mobility behaviour^29^. Yet the mechanisms linking demographic context, mobility, cultural norms, fitness consequences and evolutionary outcomes have not been investigated (Figure 1). Here, we use newly collected genome-wide data from 245 BaYaka individuals across these two regions to determine, first whether inbreeding is associated with negative fitness outcomes in a hunter-gatherer population and second whether communities use mobility strategies and cultural norms over marriage practices to mitigate inbreeding arising from differences in local demographic conditions. By tracing these connections, we provide direct evidence linking local demographic conditions, spatial behaviour, and cultural norms to genetic outcomes and reproductive fitness, shedding light on how local-scale processes shape evolutionary trajectories and emphasizing the role of culture in human adaptation.

### 1. Homozygosity results in negative fitness effects

We first tested whether homozygosity affects reproductive fitness. Using Bayesian multilevel models with a Poisson link function, we examined the relationship between F_ROH_ and the number of surviving offspring among adult and post-reproductive individuals, controlling for age, sex, and random effects of camp nested within region. We found no differences in fitness between men and women (Figure 2; Table S1-S2).

**Figure 2.**
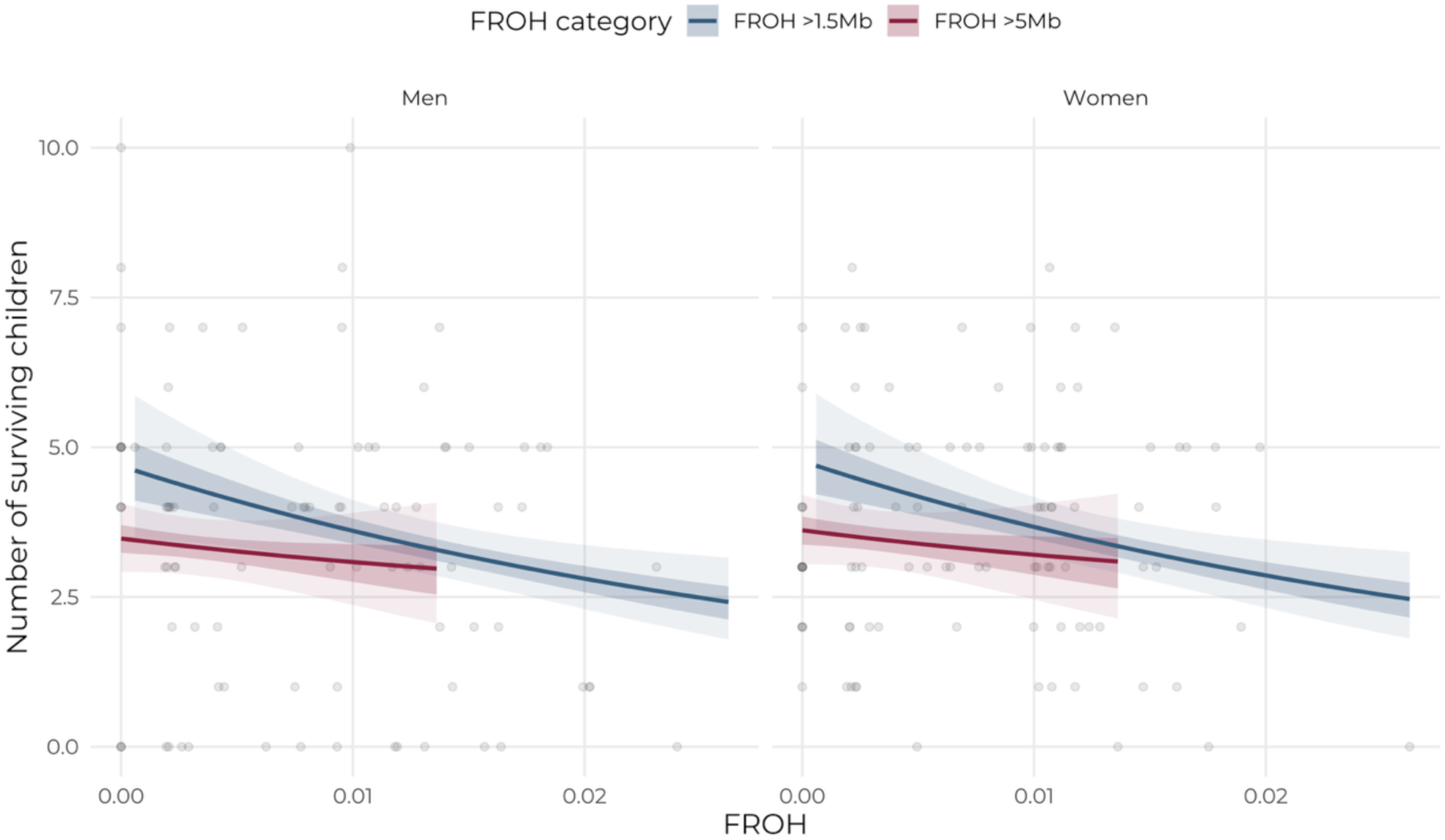
Relationship between homozygosity and reproductive fitness. Expected number of surviving offspring as a function of F_ROH_ for men (left) and women (right), calculated using two ROH length thresholds: >1.5 Mb (blue) and >5 Mb (red). Grey points represent individual observations for adults and post-reproductive individuals. Lines and shaded regions are derived from Bayesian multilevel models with a Poisson link function, including F_ROH_, sex, and age group as fixed effects, and camp nested within region as a random effect. Lines indicate posterior means, darker shaded regions represent 50% credible intervals, and lighter shaded regions represent 90% credible intervals from the posterior distribution.

However, F_ROH_ calculated from ROH >1.5 Mb (considered long ROH^30,33^) was negatively associated with reproductive fitness (Figure 2, Table S1). When considering only F_ROH_ >5 Mb, the effect was in the same direction but attenuated, likely due to the very few individuals that carried such long tracts (Figure S2-S3; Table S2). Importantly, we find direct evidence of selection against homozygosity even in the absence of close-kin marriages: among the 46 spouse pairs in our dataset, only one couple had relatedness equivalent to first cousins (inferred from a KING kinship coefficient of ≥0.06), and only four couples had relatedness equivalent to second cousins (inferred from a KING kinship coefficient of ≥0.16). This indicates that the fitness costs of homozygosity in this population arise from background relatedness rather than consanguineous unions, and that even modest elevations in inbreeding—well below levels typical of first-or second-cousin marriages—carry detectable fitness consequences.

### 2. Regional differences in demography and mobility result in differences in the clustering of relatives

Given that differences in local demographic conditions are expected to affect both genetic diversity as well as cultural and behavioural responses to maintain such genetic diversity, we used Hap-Ne-IBD^59^ to estimate recent effective population sizes (*N_e_*) on a subset of unrelated samples (excluding first- and second-degree relatives) from both regions, to determine whether both regions differed in their demographic histories. We found that hunter-gatherers (but also Bantu) in both regions experienced a strong decline in *N_e_* starting around 50 generations ago, and peaking around 10-8 generations ago (Figure 3B; Figure S4), consistent with the aftermath of the Kongo Civil war and the peak of the Atlantic slave trade in the last two decades of the 18th century^60,61^. Following this decline, *N_e_* in Minganga recovered to a slightly greater extent than in Macao.

**Figure 3.**
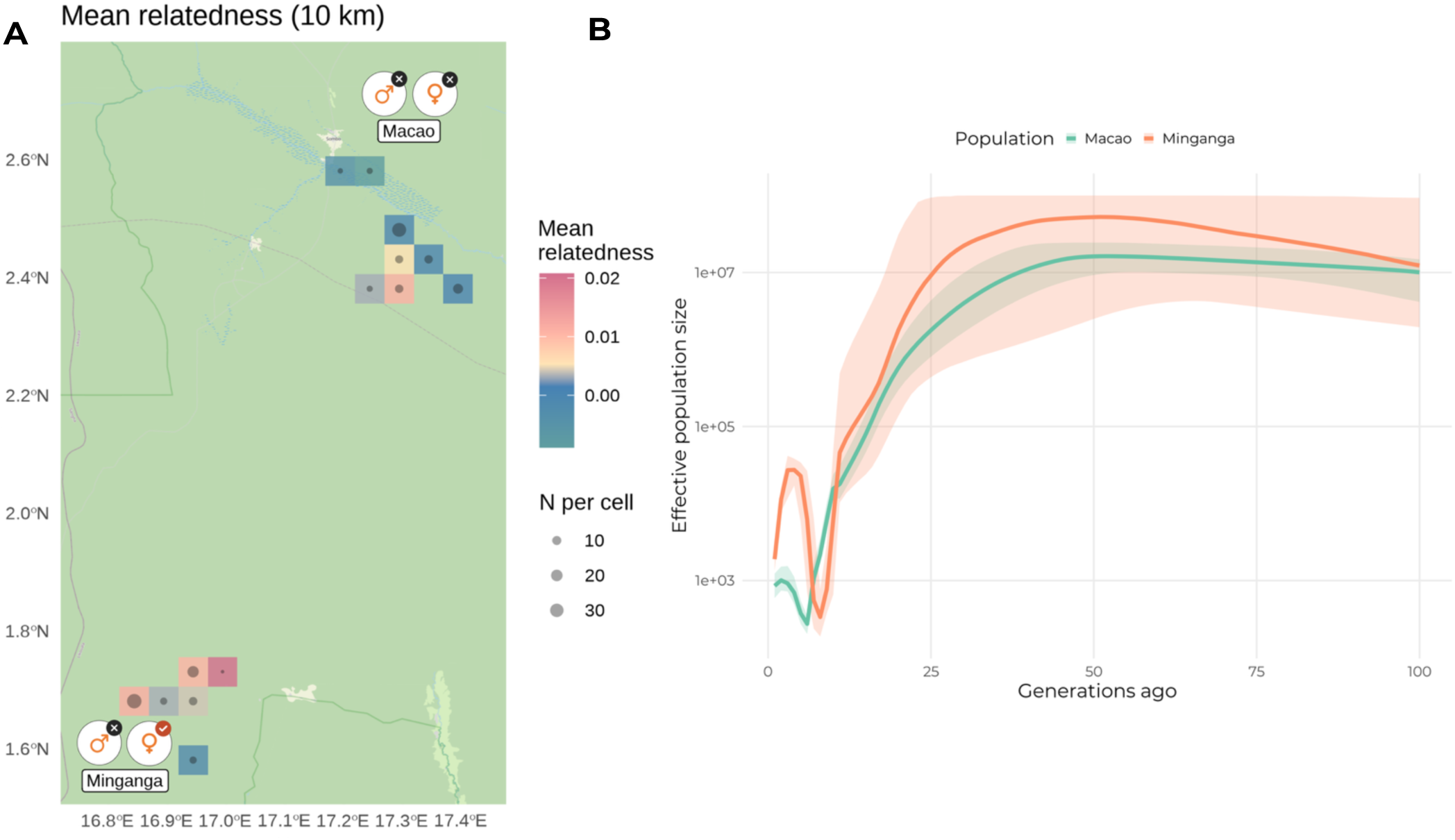
Spatial distribution of relatedness and demographic history across regions. (A) Mean relatedness of individuals to all others living within a 10km radius, calculated using KING kinship coefficients. Each square represents the mean relatedness between individuals living within 10km^2^, coloured by mean relatedness. The northern cluster corresponds to Macao and the southern cluster to Minganga. Symbols represent the exogamy norm preventing individuals from marrying others in their paternal clan (Minganga) and their paternal and maternal clans (Macao). (B) Effective population size (Ne) over time estimated using HapNe-IBD, with shaded regions indicating 95% confidence intervals. Colours indicate population: Macao (green), and Minganga (orange).

MacDonald and Hewlett^25^ predicted that regions with higher population density would show greater clustering of relatives due to reduced mate-search distances. Whilst we did not have data on population density, Bayesian multilevel models with a Gaussian link function indicated that average relatedness (mean kinship coefficient) was higher in Minganga (the region with higher *N_e_*) than in Macao (Figure 4A; Table S3-S4). This pattern is also evident when plotting mean relatedness to others within a given radius for both regions (Figure 3A; see Figure S5 for 30 km radius).

**Figure 4.**
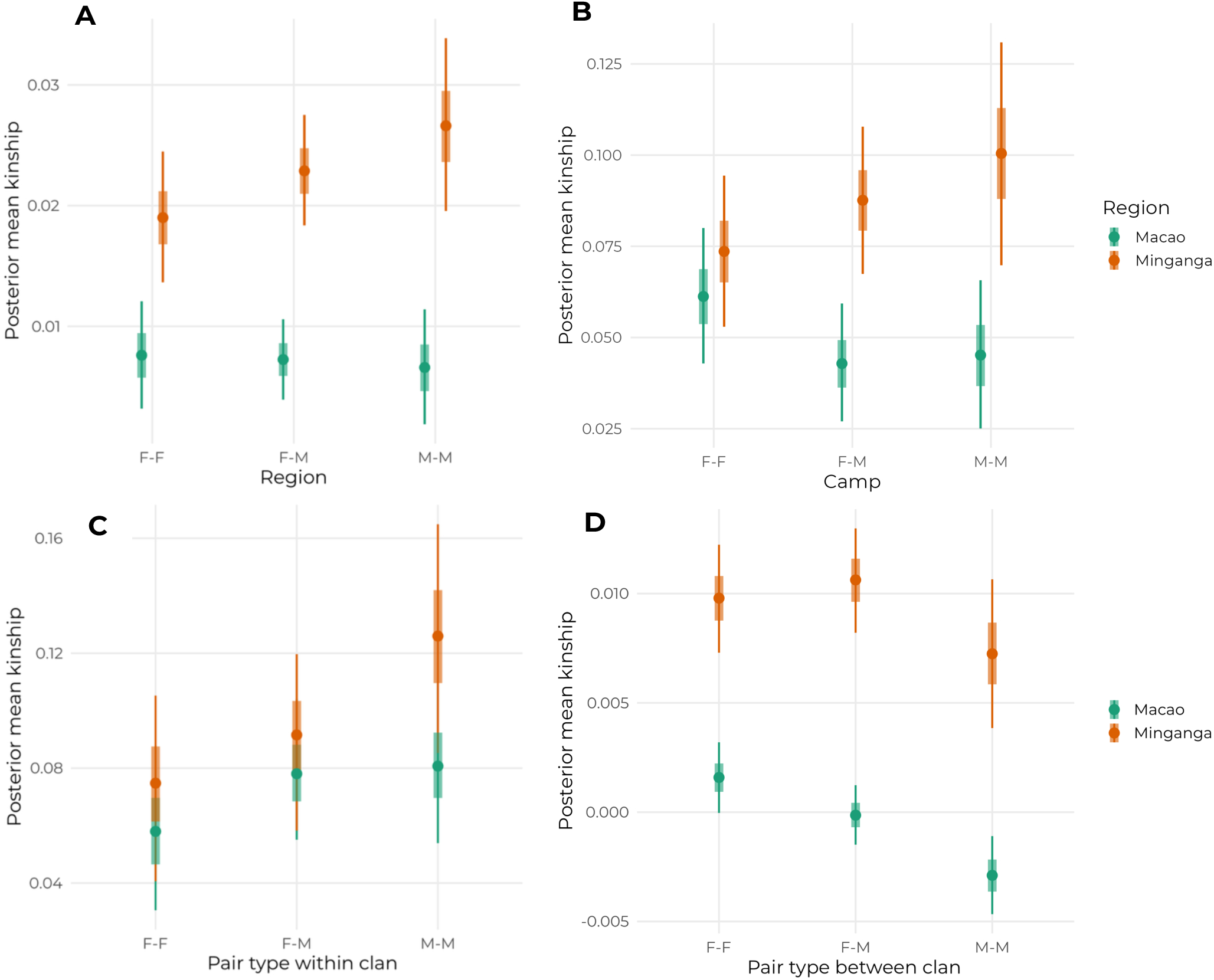
Relatedness between individuals by sex, region, and spatial scale. (A) Mean kinship between all individuals in the region by pair type (F-F: female-female; F-M: female-male; M-M: male-male) and region (Macao: green; Minganga: orange). Estimates are derived from a Bayesian multilevel model with a Gaussian link function, including pair type, region, and their interaction as fixed effects, and individual identity as a random effect. (B) Mean kinship between individuals within the same camp, estimated from a Bayesian multilevel model with pair type, region, and their interaction as fixed effects, and camp as a random effect. (C) Mean kinship between individuals within the same clan, estimated from a Bayesian multilevel model with pair type, region, and their interaction as fixed effects, and clan as a random effect. (D) Mean kinship between individuals in different clans (within regions), estimated from a Bayesian multilevel model with pair type, region, and their interaction as fixed effects, and clan as a random effect. In all panels, points indicate posterior means, thick bars represent 50% credible intervals, and thin lines represent 90% credible intervals from the posterior distribution.

However, patterns of within-camp relatedness differed between the sexes (Figure 4B). Relatedness among women was comparable across regions, whereas relatedness among men was substantially higher in Minganga (Table S5-S6; Figure S6). This suggests that in addition to differences in local *N_e_,* regional differences in post-marital residence patterns: slightly patrilocal residence in Minganga and matrilocal residence in Macao (consistent with our findings from interview data^29^), determine the clustering of relatives. Notably, interview data from Minganga indicated that women and men were equally likely to reside with their parents^29^, highlighting the complementary value of genetic data for understanding kinship structure and the spatial distribution of relatedness—patterns that may be missed by demographic surveys alone.

Finally, we examined relatedness within (Figure 4C) and between clans (Figure 4D). Between-clan relatedness was higher in Minganga than in Macao, as expected given the overall elevated relatedness in that region (Table S7-S8). However, within-clan relatedness was comparable across regions (Table S9-S10, Figure S7). Male-Female relatedness between individuals in different clans in both regions was about one order of magnitude lower than between individuals belonging to the same clan, and almost half of the average relatedness between men and women within the region. This suggests that clan exogamy norms may effectively compensate for regional differences in the background clustering of relatives.

### 3. Mate searching drives mobility for men in one region, but not the other

Differences in demographic conditions are also expected to shape mobility patterns (Figure 1). Consistent with observations from other hunter-gatherer populations^25,53^, Minganga, the region with higher historical *N_e_*, was also characterised by greater sedentism compared to Macao. This was evident both in the median and maximum half ranges of individuals from their place of residence (median distance between all the places they had visited at least once, and their current camp of residence)^29^ and in exploration ranges from birthplace estimated over individuals’ lifetimes using Bayesian multilevel models with a Gamma link function (Figure S1, Methods).

However, contrary to what was expected, mating distances were larger in Minganga (Mean posterior estimate for Minganga = 8.82km, 90% CI: [2.25-20.87] and for Macao = 3.50, 90% CI: [1.27-7.39]; Figure 5A; Table S11-S12). This finding is notable given the less stringent exogamy norm in Minganga: combined with the higher background relatedness in the region, this may compel individuals to travel further to find unrelated mates. In line with the observation that men are more related to others within their camps and clans than women are, we find that in Minganga, reproductive decision-making increases travel distance for men but not for women (Figure 5C; Table S13). This is not the case in Macao (Figure 5B). Additionally, exploration ranges from birthplace decreased in old age in Minganga (Figure S1), consistent with the prediction that travel is largely driven by the need to look for unrelated mates, which declines in later life^62^.

**Figure 5.**
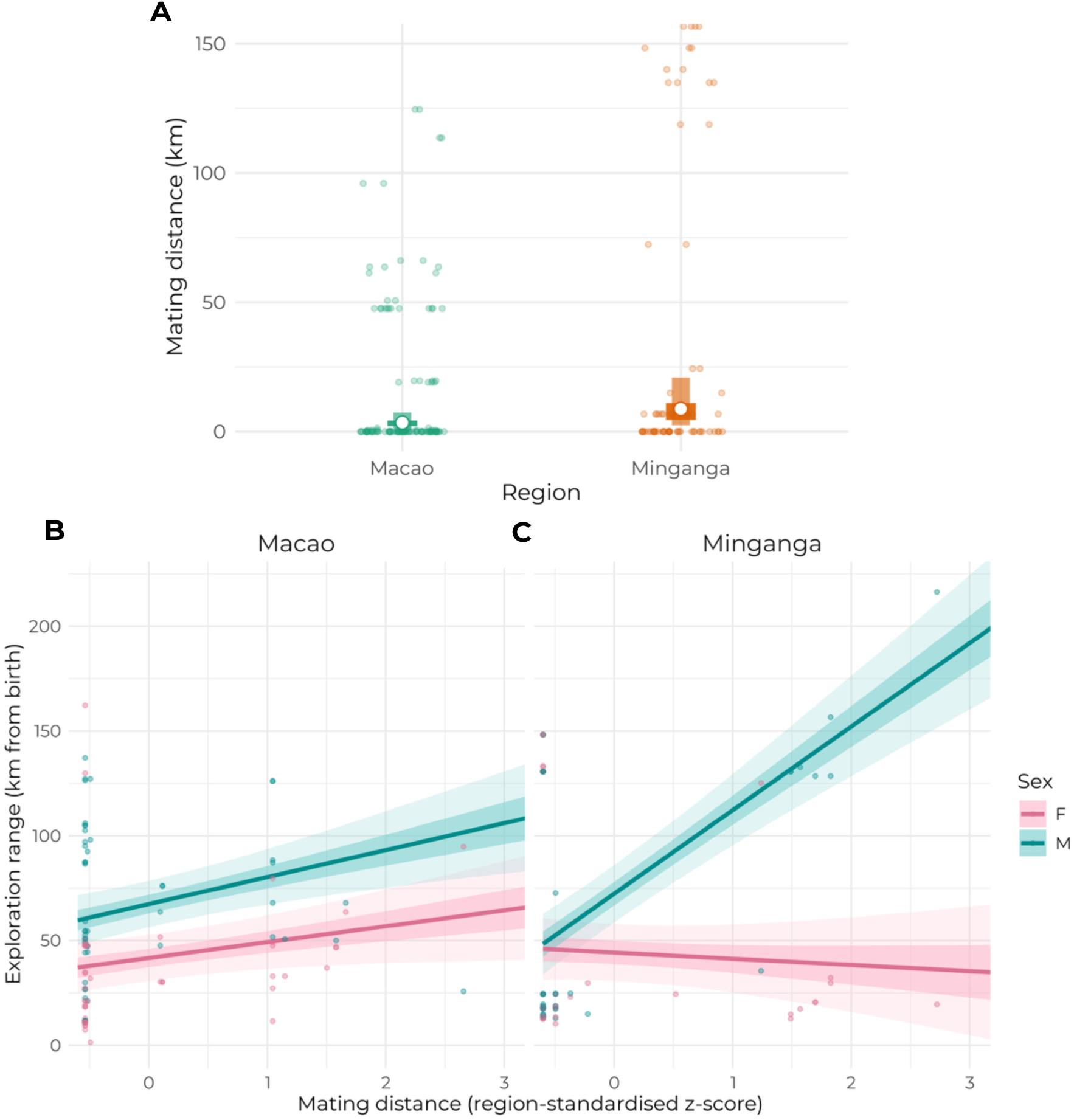
**(A) Mating distance across regions**. Mating distance, calculated as the distance between the birthplaces of spouses (in km), for couples in Macao (green) and Minganga (orange). Individual observations are shown as jittered points. Summary statistics are derived from a Bayesian multilevel model with a Gamma link function (log), with region as a fixed effect and camp nested within region as a random effect. White-filled circles indicate posterior means, thick bars represent 50% credible intervals, and lighter bars represent 90% credible intervals from the posterior distribution. (B & C) **Relationship between exploration range and mating distance by sex and region**. Exploration range (maximum distance travelled from birthplace in km) as a function of mating distance (region-standardised z-score) for females (pink) and males (blue) in Macao (B) and Minganga (C). Points represent individual observations. Lines and shaded regions are derived from a Bayesian multilevel model with a Gaussian link function, including mating distance, sex, region, and their interactions as fixed effects, age as a covariate, and camp nested within region as a random effect. Dark lines indicate posterior means, darker shaded regions represent 50% credible intervals, and lighter shaded regions represent 90% credible intervals.

### 4. Homozygosity levels are comparable across regions, but reduced by mobility

Despite the differences in demography and relatedness documented above, we found no significant differences between Macao and Minganga in either the number or total length of runs of homozygosity (ROH) in individuals’ genomes. This also held when we compared F_ROH_ between the regions, regardless of whether we used ROH > 1.5 Mb or ROH > 5 Mb (Wilcoxon rank test *p = 1;* Figure S8). Thus, contrary to predictions based on MacDonald and Hewlett^25^, differences in effective density and density of relatives (both higher in Minganga than in Macao, Figure 3B and Figure 4A), did not translate into regional differences in inbreeding levels. This suggests that the interplay between mobility and cultural norms may mitigate the potential effects of demographic differences across regions. In Minganga specifically, increased mating distances may be sufficient to reduce offspring inbreeding even under a more relaxed exogamy norm that prohibits marriage only within the paternal clan.

To test whether mobility directly affects offspring inbreeding load, we identified individuals for whom both parents were present in the dataset and examined the relationship between parental mating distance and offspring ROH using Bayesian multilevel models with a Beta link function. Previous work has suggested that F_ROH_ > 1.5 Mb correlates with pedigree-based inbreeding coefficients^30^, though more recent studies indicate that across diverse samples, pedigree-based inbreeding correlates well with F_ROH_ calculated from ROH > 5 Mb but not from shorter segments^33,63^. We therefore present results using both thresholds.

Among individuals with both parents in the dataset (a sample which mostly comprised children and adolescents – as these age groups were most likely to have both parents alive, and therefore included in our genomic and ethnographic datasets), offspring in Minganga showed higher inbreeding levels than those in Macao when considering both F_ROH_ > 1.5 Mb and F_ROH_ > 5 Mb (Figure 6; Table S14-S15). This suggests that inbreeding has increased in recent generations in Minganga - a pattern not evident in the full sample. Crucially, parental mating distance (the distance between parents’ birthplaces) negatively predicted very long ROH in offspring (i.e. F_ROH_ > 5 Mb; Figure 6B). We tested whether this effect differed between regions by including an interaction between region and parental mating distance, but in all cases models without the interaction were better supported (Table S16), indicating that the protective effect of mobility on offspring inbreeding operates similarly across both regions.

**Figure 6.**
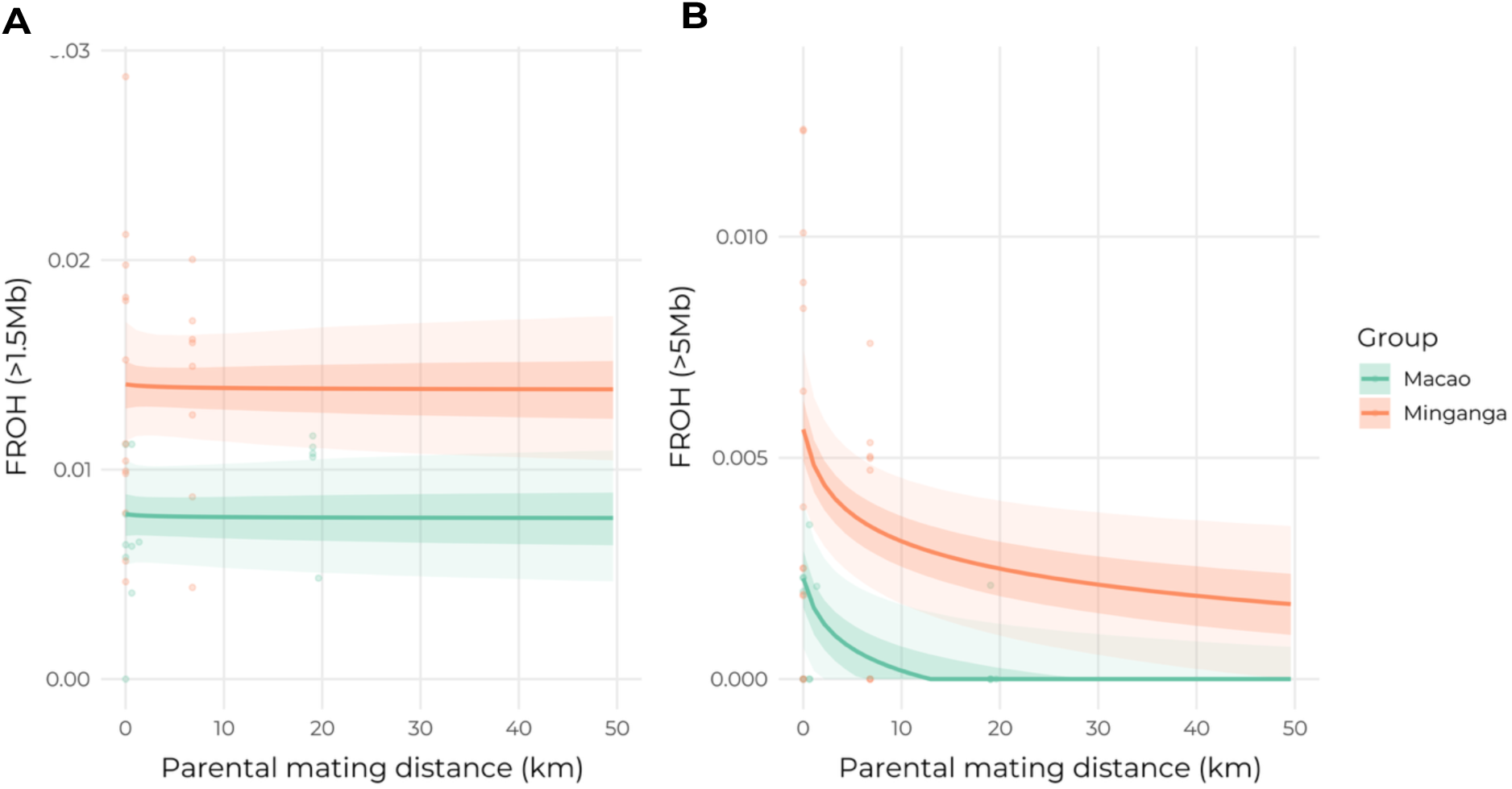
Relationship between parental mating distance and offspring inbreeding. Fraction of the genome in runs of homozygosity (F_ROH_) as a function of parental mating distance (distance between parents’ birthplaces in km) for offspring in Macao (green) and Minganga (orange). (A) F_ROH_ calculated from ROH > 1.5 Mb; (B) FROH calculated from ROH > 5 Mb. Points represent individual observations. Lines and shaded regions are derived from Bayesian multilevel models with a Beta link function, including log-transformed parental mating distance and region as fixed effects, and camp as a random effect. Lines indicate posterior means, darker shaded regions represent 50% credible intervals, and lighter shaded regions represent 90% credible intervals from the posterior distribution.

## Discussion

Elevated homozygosity is associated with reduced fitness across a wide range of species, implying strong selection against inbreeding, yet its fitness consequences remain underexplored in humans. Here, we show that even in a hunter-gatherer context characterised by a near absence of close-kin marriages, increased homozygosity is associated with reduced reproductive success. Importantly, these effects are driven by runs of homozygosity longer than 1.5 Mb, which reflect recent background relatedness rather than explicit consanguineous unions. Together, these findings demonstrate that maintaining genetic diversity is an evolutionarily consequential target of selection.

Given such selection pressure, we would expect human populations to develop a suite of cultural and behavioural strategies to maintain genetic diversity under different local demographic, social and ecological conditions. Yet genetic and ethnographic evidence for such strategies have largely remained disconnected. By integrating genomic, demographic, mobility, and cultural data from two Central African hunter-gatherer populations in the northern Republic of Congo (Macao and Minganga), we show that despite regional differences in historical effective population sizes and background relatedness, overall levels of homozygosity are remarkably low and highly similar. These shared genetic outcomes are achieved through distinct combinations of mobility patterns and clan exogamy norms, illuminating how culture and behaviour interact to shape evolutionary processes (Figure 1).

In Minganga, despite a higher clustering of relatives, greater sedentism and a more relaxed exogamy norm, men’s greater travelling distance to find mates might buffer offspring from increased homozygosity, even under conditions of elevated background relatedness. On the other hand, in Macao, reduced sedentism (increased mobility over the lifetime) and strict exogamy norms preventing individuals from marrying those from both their maternal and paternal clans appear to play a central role in maintaining genetic diversity. These mechanisms reduce the need for men to travel very long distances to find spouses by keeping a low density of relatives around their camps of residence, despite the lower effective population size observed in this Macao. In both regions, the existing clan exogamy norms ensure that spouses are less related than randomly drawn individuals from the population or within clans.

This result challenges a common assumption in spatial population genetics, namely that local concentrations of relatives straightforwardly reflect local population size or density^54,64–66^. In this hunter-gatherer groups, social organisation and exogamy rules decouple local census size from local genetic structure, breaking the expected association between effective population size and relatedness (Figure 2A). This highlights the need to explicitly incorporate cultural institutions, practices and social organization into models of human spatial genetic structure.

Our results indicate that in Minganga, but not in Macao, sex differences in spatial behaviour are at least partly driven by mating effort (see Padilla-Iglesias et al.^29^ for drivers of sex differences in mobility in Macao). This is evidenced by the positive association between lifetime exploration range and mating distance for men (but not for women) and by the decline in exploration range from adulthood to old age. This pattern mirrors findings across diverse societies, including the Yucatec Maya in Mexico and the Twe and Himba in Namibia, where sex differences in mobility emerge most strongly during young adulthood, when mate search is most relevant, and decline later in life^62,67^. In these populations, men who travel more also tend to achieve greater mating success^68^. Importantly, our results suggest that such sex differences in reproductively-driven spatial behaviour are not universal, but contingent on local demographic conditions.

Our findings are in line with earlier work showing that sedentarisation is often associated with increased mating distances^53,69^. In Minganga, higher sedentism is accompanied by longer mating distances, likely because elevated background relatedness necessitates greater mate-search effort under relaxed exogamy rules.

Our findings have broader implications for how spatial genetic processes are studied in humans. Much of spatial population genomics has focused on “dispersal distance”, often defined as the distance between parents’ and offspring’s birthplaces, as the key parameter governing genetic outcomes^54^. However, in this group, individuals may travel extensively during their lifetimes yet return to the same locations to give birth^29,45,70–72^. As a result, dispersal distance alone may severely underestimate the extent to which populations are socially and genetically connected. To understand the drivers of human genetic evolution, more studies should move beyond static measures of dispersal and incorporate lifetime mobility, mate-search behaviour, and post-marital residence patterns.

We provide evidence that long runs of homozygosity are associated with reduced reproductive success in a population characterised by a near absence of close-kin marriages. Consistent with this, fitness effects were driven by F_ROH_ calculated from segments >1.5 Mb, which capture recent background relatedness rather than consanguineous marriages^33^. Earlier work on mobility among the BaYaka of Macao and Minganga found a link between mobility and reproductive success^29^. Here we provide a plausible mechanism linking the two by showing that mobility reduces inbreeding load, and in turn that reduced inbreeding load has positive fitness consequences – directly linking spatial behaviour with evolutionarily-relevant outcomes.

In conclusion, by directly investigating the relationship between demography, cultural norms, spatial behaviour, genetics, and fitness outcomes, we provide a mechanistic account of how local-scale processes shape human evolutionary trajectories. Our findings reveal how culture is an active component of human adaptation^73^, modulating mobility, regulating reproduction, and ultimately influencing genetic diversity and reproductive success. In doing so, they help explain the resilience of the hunter-gatherer foraging niche despite demographic and climatic shocks.

## Materials and methods

### Study population and data collection

The BaYaka are a hunter-gatherer population living in the rainforests of the Congo Basin. They belong to a broader group of hunter-gatherers collectively known as Central African hunter-gatherers ^74,75^ or rainforest hunter-gatherers^76,77^ that have inhabited the rainforests of West and Central Africa since at least the last interglacial ^26,28^ (See Text S1 in Padilla-Iglesias et al.^29^ for extensive ethnographic information).

For the present study, we surveyed two different regions (broad research areas) in the Likouala department of the Republic of Congo. We call them after the major hunter-gatherer settlement in each (Macao and Minganga, respectively, as have other scholars^71,78^ even if we did not survey the villages themselves). Both regions were located within forest environments and around 110km apart from one another. Hunter-gatherer populations in the region vary in residential mobility, dialect (of the same language), and frequency of foraging as well as seasonal work in agriculture for neighbouring farming populations. They also differ in their access to roads and motorized means of transport, local population density and cultural norms for family organization.

Data collection was conducted between March and June 2023. During field visits to camps in each of the two study regions, genetic, demographic and mobility data was collected for all adolescent and adult individuals who were present, provided informed consent, and agreed to participate after being informed of the study’s aims and procedures and saliva samples were collected from children whose parents’ provided informed consent in their behalf. Participants sometimes volunteered for providing certain types of data and not others. In addition, For each region, we also collected saliva samples from N=20 consenting Bantu-speaking agriculturalists living in the vicinity of the BaYaka. Saliva samples were collected using Oragene-OG600 DNA collection kits following the protocols specified by the manufacturer and stored at room temperature. DNA amples were genotyped with the H3Africa genotyping array^79^, which is specifically designed for sampling genomic variation within Africa. Quality control (QC) was performed and succeeded for N=287 samples (N=33 of them which were from Bantu-speaking agriculturalists).

Demographic and mobility data for the BaYaka were collected and coded using the exact same protocol detailed in Padilla-Iglesias et al.^29^.

### Genetic data filtering and processing

We used the R package *tidypopgen* (version 0.4.2)^80^ for all subsequent genetic filtering and analyses unless otherwise stated. We began with autosomal genotype data after removal of duplicate markers. To minimize the influence of close kinship on IBD-based demographic inference, individuals related more closely than second degree were excluded using KING estimates (kinship coefficient > 0.088).

Variants and individuals were filtered using the following criteria: individual missingness < 20%, per-locus missingness < 20%, and minor allele frequency (MAF) > 1%. All filtering steps were applied iteratively after sample removal to ensure consistency of the final dataset. We then grouped individuals by population and filtered out loci that were not in Hardy-Weinberg equilibrium (p>0.001). After such filtering was performed, we were left with 1,943,877 autosomal SNPs and N=273 individuals.

In addition, prior to performing the ADMIXTURE^81^ analysis, LD-based pruning was applied to remove SNPs in high linkage disequilibrium using an 𝑟^2^ threshold of 0.1 within 1 Mb genomic windows. This resulted in a dataset of 211,147 autosomal SNPs and N=273 individuals.

We further applied Principal Component Analyses to visually inspect the data. To place our samples within a broader African context for ADMIXTURE analyses and analyses of ROH we merged the data with that from studies including other African populations^5,82–84^, and applied the same filters as specified above. For PCA and ROH analyses, this left us with N=914 individuals and 379,204 autosomal SNPs. For ADMIXTURE analyses, we further subsetted the merged dataset to individuals from Western-Central African populations, and after LD-based pruning, this left us with N=810 individuals and 138,940 autosomal SNPs.

### Detection of runs of homozygosity and F_ROH_ calculations

Runs of homozygosity (ROH) were identified using a sliding-window approach. Analyses were performed separately for each population. To minimize spurious ROH calls we required each ROH to contain at least 30 SNPs, span a minimum physical length of 500 kb, and meet a minimum SNP density threshold of one SNP per 30 kb allowing a maximum of one heterozygous site and up to five missing genotypes per window^85,86^. The resulting ROH set was used for all downstream analyses. For the analyses reported in the main manuscript, we focused on ROH >1.5Mb and ROH >5Mb in length, as different studies have claimed that these categories correlate strongest with pedigree consanguinity^30,33,43,63^.

Then, we calculated the fraction of individuals’ genomes in runs of homozygosity (F_ROH_^31,43,85^) as the total length of ROH longer than 1.5 Mb and 5Mb divided by the total length of the autosomal genome reliably covered by the genotyping array (2,713,905,338 base pairs).

### Phasing of genotypes

Filtered genotype data were converted to VCF format and split by chromosome. Phasing was performed separately for each population using Beagle (version 4.2)^87^, with genetic maps corresponding to GRCh37 and centromeric and telomeric regions removed. Phasing was conducted chromosome by chromosome using a sliding window of 20 cM and a random seed to ensure reproducibility (as recommended in the software manual). Phased chromosomes were subsequently recombined into population-specific genome-wide phased VCFs and reformatted to ensure compatibility with downstream analyses.

To meet the requirements of HapNe-IBD, phased VCFs were further split into chromosome arms using custom scripts, allowing IBD length distributions to be analysed independently across arms.

### Detection of IBD segments and estimation of effective population size through time

IBD segments were detected from phased data using Hap-IBD^88^ (version 1.0). We retained IBD segments with a minimum genetic length of 4 cM, corresponding to recent shared ancestry^43^. To reduce fragmentation of long IBD tracts, adjacent segments were merged if separated by gaps shorter than 0.6 cM, allowing at most one discordant homozygous genotype, as recommended in the software manual.

Effective population sizes (*N_e_*) for the past 100 generations were inferred using HapNe-IBD, which models the distribution of IBD segment lengths to estimate *N_e_* across time. For each population, IBD length histograms were constructed across chromosome arms and used as input to the HapNe-IBD framework.

### Ethical approval and consent

Approval for this research was obtained from the Ethics Commission (Ethikkommission) of the University of Zurich (Permit Nr. 20.2.8), the Canton of Zürich, the Ethics Committee at the University of Cambridge (Permit HBREC.2025.21) as well as by the Presidency of Research and Cooperation from the Marien Ngouabi University, the Ministry of Justice, Human Rights and promotion of indigenous peoples of the Republic of Congo, and the Ministry of the Environment and Sustainable Development of the Republic of Congo. All participants provided informed consent after receiving a comprehensive explanation of the study’s objectives, the data to be collected, and the broader research goals. This information was presented orally, with questions answered to ensure that consent was fully informed.

### Ethics and inclusion statement

This research was conceptualised from its inception in collaboration with researchers from Marien Ngouabi University, Republic of Congo (co-authors D. Nganga, E. Amboulou, and S.I. Averti), who participated in study design, data collection, data processing, and manuscript preparation. All authors have equal rights to the content of this manuscript. The study was conducted in collaboration with the Ministry of Justice, Human Rights and Promotion of Indigenous Peoples of the Republic of Congo, and the Ministry of the Environment and Sustainable Development of the Republic of Congo.

Locally relevant issues, beliefs and cultural practices were taken into consideration when designing research questions and methodologies and were refined through pilot visits to all study communities prior to data collection. Community assemblies were organised in each location prior to data collection to explain the research objectives, methods, and the roles of the international team. Community members were invited to express reservations, interests, and suggestions, and research methodologies were adapted accordingly. All communities responded positively to participation. In addition, informed consent was obtained individually from all participants (see *Ethical approval and consent*).

### Statistical approach

We used Bayesian inference for all statistical analyses, which allows for comprehensive interpretation of posterior distributions and provides interval estimates that reveal the range of plausible values for each parameter. For all models, we report the 50% and 90% credible intervals (CI), which represent the narrowest interval containing 50% and 90% of the posterior probability for each parameter^89^.

We chose priors that were regularising in order to prevent the model from overfitting the data^89^, as well as being biologically motivated. In addition, predictive checks were performed to make sure all priors were consistent with characteristics of each parameter. We further fitted alternative parameterizations for all models, to confirm that the results presented below were qualitatively robust to changes in priors. Parameter estimation was achieved with the *brms* (v2.22.0) R package^90^. Models were fitted using four Markov chain Monte Carlo chains, each run for 5,000 iterations with a warm-up of 1,000 iterations, and convergence was achieved for all models, suggested by a high effective number of samples and R^ estimates of 1.00. Trace plots were also inspected to make sure chains converged to the same target distributions and posterior predictions were compared to the raw data to confirm the match between model predictions and the descriptive summaries of the samples.

### Pairwise relatedness across geographic and social scales

To characterise fine-scale genetic structure and assess whether pairwise relatedness (between dyads) varied by sex composition and region across different social and geographic scales, we modelled pairwise kinship coefficients using a series of Bayesian mixed-effects models. All models shared the same fixed-effects structure but differed in the level at which non-independence among dyads was modelled, allowing relatedness to be examined across nested social units: regions, camps, clans, and between clans.

Pairwise kinship coefficients were calculated for all pairs of individuals using *tidypopgen*^80^. Dyads were annotated with region and pair type, where pair type was coded as a three-level categorical variable based on the sex of the two individuals (F–F, F–M, M–M).

Region-wide relatedness was first estimated using all dyads within regions while accounting for repeated appearance of individuals across dyads by including crossed random intercepts for both members of each pair:

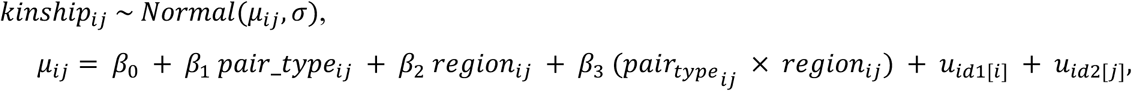

where u_id1[i]_ and u_id2[j]_ are random intercepts for the two individuals in each dyad.

For this model, priors were specified as follows: the intercept was assigned a *Normal(0.005, 0.04)* prior, reflecting low baseline relatedness across the region; regression coefficients were assigned *Normal(0, 0.04)* priors; standard deviations of the individual-level random effects were assigned *Student-t(3, 0, 0.03)* priors; and the residual standard deviation was assigned a *Student-t(3, 0, 0.05)* prior.

We next restricted the dataset to dyads where both individuals belonged to the same camp and region, and fitted an analogous model with a camp-level random intercept:

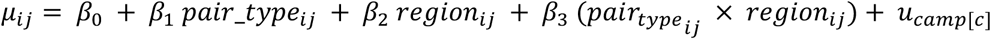

For the within-camp model, priors were identical to the region-wide model except that the intercept was centred on higher expected relatedness *(Normal(0.05, 0.04))*, reflecting the a priori expectation that individuals within the same camp are more related on average.

We then created a dataset including dyads in which both individuals belonged to the same clan, and fitted the same fixed-effects structure with a clan-level random intercept:

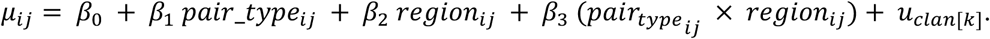

Priors for the within-clan model were chosen to reflect higher expected relatedness and lower variance among clans: the intercept was assigned a *Normal(0.05, 0.04)* prior; regression coefficients were assigned *Normal(0, 0.04)* priors; the standard deviation of the clan-level random effect was assigned a *Student-t(3, 0, 0.02)* prior; and the residual standard deviation was assigned a *Student-t(3, 0, 0.03)* prior.

Finally, to quantify relatedness between clans, we modelled dyads belonging to different clans using a random intercept for clan pairs:

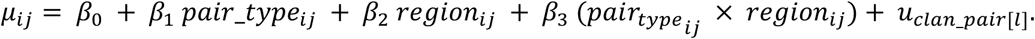

For the between-clan model, priors reflected low expected relatedness between clans: the intercept was assigned a *Normal(0.002, 0.005)* prior; regression coefficients were assigned *Normal(0, 0.01)* priors; the standard deviation of the clan-pair random effect was assigned a *Student-t(3, 0, 0.01)* prior; and the residual standard deviation was assigned a *Student-t(3, 0, 0.02)* prior.

### Regional differences in mating distances

To test whether parental mating distance varied between regions, we modelled mating distance (km) as a function of region while accounting for clustering by residence camp. Mating distance was defined as the geographic distance between spouses’ birthplaces^25^. Because some observations were equal to zero, we added a small offset prior to modelling (0 km replaced with 0.001 km) to ensure compatibility with positive-support distributions^89^.

We fitted Bayesian generalized linear mixed-effects models. Distances were modelled with a Gamma likelihood to reflect their strictly positive, right-skewed distribution. The primary model used a log link:

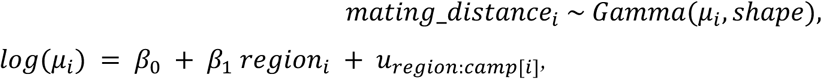

where μ_i_ is the expected mating distance for individual *i, region_i_* is a categorical predictor (Macao vs Minganga), and *u_region:camp[i]_* is a random intercept for residence camp nested within region, assumed to follow a normal distribution with mean 0 and variance *σ²_region:camp_.* The Gamma shape parameter was estimated from the data.

On the log scale, the intercept was assigned a *Normal(log(10), 0.5)* prior. The region effect was assigned a *Normal(0.5, 1)* prior. Standard deviations of random effects were assigned *Student-t(3, 0, 0.5)* priors. The Gamma shape parameter was assigned a *Lognormal(log(3), 0.5)* prior.

### Relationship between mating distance and exploration range

To determine whether the search for mates drives hunter-gatherer spatial ranges, we examined whether individuals’ mating distance predicted their exploration ranges.

Exploration range was measured in kilometres as the median distance between an individual’s birthplace all the places they had visited at least once (see^29^ for details on how these data were collected). Because parental mating distances were right-skewed and included zeros, we applied a log transformation defined at zero.

We fitted Bayesian generalized linear mixed-effects models. Exploration range on the original scale was analysed assuming a Gaussian likelihood with identity link:

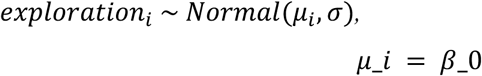

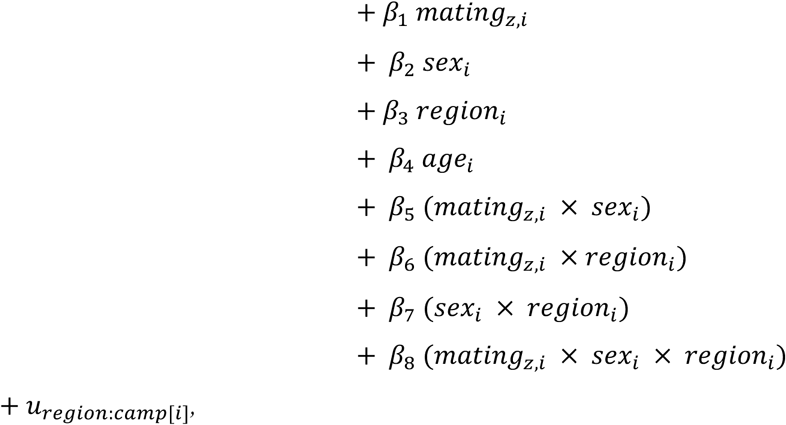

where *mating_z,i_* denotes parental mating distance standardised within region, and *u_region:camp[i]_* is a random intercept for camp nested within region, assumed to follow a normal distribution with mean 0 and variance *σ²_region:camp_*.

The intercept was assigned a *Normal(50, 30)* prior, corresponding to plausible values of exploration range in kilometres. Regression coefficients were assigned *Normal(0, 30)* priors. Standard deviations of random effects and the residual standard deviation were assigned *Student-t(3, 0, 30)* priors.

### Relationship between mating distance and offspring homozygosity

To test whether greater parental mobility is associated with reduced genomic homozygosity in offspring, we modeled individuals’ fraction of the autosomal genome contained in runs of homozygosity (F_ROH_) as a function of parental mating distance. Clustering at the camp level was explicitly accounted for by including random effects for camp in all models.

Because parental mating distances were right-skewed and included zeros, we applied a log transformation defined at zero. Similarly, because F_ROH_ is a continuous proportion that can include exact zeros, we modeled FROH using beta regression. To satisfy the requirement that beta-distributed outcomes lie strictly within the open interval (0,1), we applied the Smithson and Verkuilen^91^ adjustment:

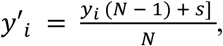

where 𝑦*_i_* is the observed F_ROH_ value, *N* is the sample size, and *s* is a constant between 0 and 1. We set *s* = 0.5, which corresponds to mild shrinkage away from the boundaries and can be interpreted as incorporating weak prior information. This transformation was applied separately to F_ROH_ > 1.5 Mb and F_ROH_ > 5 Mb.

We fitted Bayesian generalized linear mixed-effects models with a beta likelihood with a logit link for the mean. For individual i:

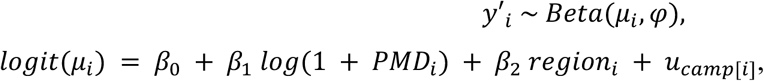

where *PMD_i_* is parental mating distance in kilometers, region (Macao or Minganga), and u_camp[i]_ is a random intercept for camp, assumed to follow a normal distribution with mean 0 and variance σ^2^_camp_. The beta precision parameter φ was modelled as constant across observations.

On the logit scale, the intercept was assigned a normal prior centred on low background homozygosity *Normal(−4, 1)*. Regression coefficients for log-transformed parental mating distance and group were assigned *Normal(0, 0.5)* priors, constraining effect sizes to biologically plausible ranges while allowing sufficient flexibility. The intercept for the precision parameter φ was assigned a *Normal(7.5, 1)* prior.

We compared models with and without an interaction between parental mating distance and region using leave-one-out cross-validation (LOO-CV^92^).

### Relationship between homozygosity and reproductive success

To test whether parental genomic homozygosity predicts reproductive success, we modelled the number of surviving children per individual at the time of data collection as a function of homozygosity. We considered two measures of genomic homozygosity: F_ROH_ calculated from runs of homozygosity longer than 1.5 Mb and FROH calculated from runs longer than 5 Mb. To aid interpretability and improve our Bayesian models’ sampling efficiency, F_ROH_ predictors were standardised prior to analysis.

We fitted Bayesian generalized linear mixed-effects models with a Poisson likelihood with a log link. For individual *i*:

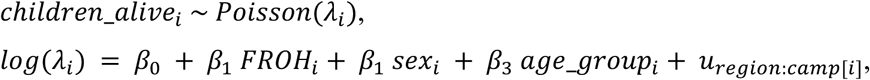

where 𝑠𝑒𝑥*_i_* and 𝑎𝑔𝑒_𝑔𝑟𝑜𝑢𝑝*_i_* were included as categorical fixed effects, and *u_region:camp[i]_* is a random intercept for camp nested within region, assumed to follow a normal distribution with mean 0 and variance σ²_region:camp_. Separate models were fitted for F_ROH_ >1.5 Mb and F_ROH_ >5 Mb.

The intercept was assigned a *Normal(−1, 1)* prior on the log scale, corresponding to low expected numbers of surviving offspring. Regression coefficients were assigned *Normal(0, 0.5*) priors. Standard deviations of random effects were assigned *Student-t(3, 0, 0.5)* priors.

## Acknowledgements

We are grateful to the BaYaka and Bantu individuals and communities for their participation in this study, and for guiding us through our research questions and methodologies. We thank the iGE3 Genomics Platform for support with genetic data processing, and H3Africa for genotyping and technical support. We also acknowledge the Centre for Genomic Regulation (CEGEN, Santiago) for assistance with DNA extractions. We also wish to thank Pier Palamara for guidance on running HapNe-IBD. We thank Emmanuel College, University of Cambridge, and the University of Zurich for institutional and financial support. This work was also funded in part by the Leakey Foundation and the A. H. Schultz Foundation. Last, we thank Sarah Tishkoff and Etienne Patin for access to genomic datasets.

## Data availability

Raw genome-wide SNP data were obtained from the datasets reported in Perry et al., 2014; Patin et al. 2014; 2017; Jarvis et al. 2012 and are available in the public servers EGA (accession numbers EGAS00001002078 and EGAC00001000139) and dbGaP (accession numbers phs000449.v2.p1 and phs001780.v1.p1). Newly generated data will be uploaded to EGA upon acceptance of the manuscript.

## Code availability

All the code required to reproduce the results in this manuscript is available in the following GitHub repository: https://github.com/EvolEcolGroup/congo_genotype_analyses

